# Positive impact of Type-A and -C response regulator genes on growth and panicle architecture in rice

**DOI:** 10.1101/2024.10.28.620749

**Authors:** Chenyu Rong, Renren Zhang, Jing Xie, Jieru Li, Tiantian Yan, Ziyu Liu, Yuexin Liu, Ruihan Xu, Xi’an Shi, Xuebin Zhao, Jiali Song, Yayi Meng, Zhongyuan Chang, Yanfeng Ding, Chengqiang Ding

## Abstract

Cytokinin signal transduction occurs through a "two-component system." Type-A and -C response regulators (RRs) are groups of proteins of similar structures constituting significant components of cytokinin signal transduction. In rice, 13 (Type-A) and two (Type–C) *RRs* have been identified to date; however, their functions remain partially known. In this study, we examined the expression patterns of Type-A and Type-C RRs in rice using RNA-Seq and confirmed their functions by constructing mutants of the 15 genes with CRISPR/Cas9. Almost all Type-A *RRs* played positive roles in the development of secondary branches and secondary spikelets, particularly *RR2* and *RR4*. Notably, *rr1 rr2* and *rr8 rr12 rr13* higher-order mutants displayed small panicle sizes and decreased plant height. Additionally, both Type-C *RRs* played positive roles in regulating heading date. RNA-seq revealed several genes with significantly altered expression in the *rr2* and *rr4* mutants, with almost half of the differentially expressed genes (DEGs) overlapping between the two mutants. Many of the DEGs were associated with the cytokinin and abscisic acid pathways. Our findings provide new insights into the functions of Type-A and -C *RRs* in rice growth and may serve as a foundation for future studies focusing on cytokinin signaling.

## Introduction

Cytokinins are crucial in cell division, root and shoot growth, senescence, and communication between plants and their environment (Mok and Mok, 2001; Zurcher and Muller, 2016). Cytokinin functions are influenced by their biosynthesis, transportation, biodegradation, and signal transduction. Most studies have shown that cytokinin signal transduction occurs through a "two-component system." Several components, such as histidine kinases (HKs), authentic histidine phosphotransfer proteins (AHPs), and Type-A, -B, and -C response regulators (RRs), participate in this process. Signal transduction is a complex process that involves various mechanisms. Despite the extensive research, many aspects of cytokinin-mediated signal transduction remain unclear.

Cytokinin signal transduction starts at the endoplasmic reticulum and plasma membrane (Romanov *et al*., 2018). Cytokinin signal transduction primarily begins with cytokinin sensor HKs, which bind cytokinins at the CHASE domain and transmit the signal by transferring a phosphate group from the histidine (His) in the sensing domain to the aspartic acid (Asp) in the receiver domain (Burr *et al*., 2020; Zurcher and Muller, 2016). Subsequently, the His in the AHPs accepts phosphate from the HKs. Finally, AHPs transfer the phosphate signal to the nucleus via the Asp on Type B RR proteins. Type B RR proteins are transcription factors that, upon activation by a phosphate signal, regulate the expression of downstream genes across the genome (Meng *et al*., 2017; Zubo *et al*., 2017). A recent study confirmed that unphosphorylated Type B RRs adopt a closed conformation, positioning the DNA recognition helix in a manner unfavorable for DNA binding. When the receiver domain of the Type-B RR protein receives phosphate, a structural rearrangement occurs, allowing Type-B RRs to demonstrate DNA-binding activity (Zhou *et al*., 2024). In addition, Type-A and -C RRs are considered inhibitors of cytokinin signal transduction, as they may disturb phosphate transfer from AHP to Type-B RR protein.

RRs are crucial components of cytokinin signaling and can be classified into three types based on their sequences and domain structures. Type B RRs have long amino acid sequences and comprise three domains: an N-terminal receiver domain that accepts phosphate transfer, a central Myb-like DNA-binding domain, and a C-terminal transactivation domain. Type A RRs have short amino acid sequences that contain a receiver domain and short N- and C-terminal extensions. Type C RRs have a structure similar to that of Type A RRs, and the two types are distinguished through phylogenetic analysis (Ito and Kurata, 2006; Tsai *et al*., 2012). In the rice genome, there are thirteen type-A *RRs* (*RR1*-*RR13*) and two type-C RRs (*RR41* and *RR42*), and type-A RRs can be divided into four subfamilies (A-II, A-IV, A-V, and A-VI) (Tsai *et al*., 2012).

In Arabidopsis and rice, the expression of most Type-A *RRs* is rapidly upregulated, primarily by exogenous cytokinins (Jain *et al*., 2006; Tsai *et al*., 2012). In contrast, the expression of Type-B and -C *RRs* do not respond to exogenous cytokinins. Other plant hormones, such as auxins, abscisic acid (ABA), and strigolactone, can also influence the expression of Type-A *RRs* in rice and Arabidopsis (Duan *et al*., 2019; Tsai *et al*., 2012; Zwack and Rashotte, 2015), suggesting that Type-A *RRs* also serve as communication bridges between cytokinins and other plant hormones. Furthermore, some Type A *RRs* respond to abiotic stress in Arabidopsis and rice (Wang *et al*., 2019; Zwack and Rashotte, 2015).

As both gene groups seem to inhibit cytokinin signal transduction, researchers have focused on their cytokinin sensitivity. In Arabidopsis, the quadruple mutant *arr3 arr4 arr5 arr6* exhibited a normal root phenotype but was more sensitive to cytokinins than the wild type (WT) was (To *et al*., 2007). Further studies also confirmed that substitution at the Asp site can cause a reduced inhibitory effect on cytokinin signaling. In contrast, some Type-A *ARR*-overexpressing lines, such as *ARR3-ox* and *ARR16-ox*, have longer primary roots, greater lateral root density, and less sensitivity to exogenous cytokinins (Ren *et al*., 2009). Similarly, in rice, Type-A *RRs* participate in cytokinin signaling, and *OsRR3* and *OsRR5* overexpression causes reduced cytokinin sensitivity (Cheng *et al*., 2010). In addition to cytokinin signal transduction, Type-A and -C *RRs* influence plant growth and development. In Arabidopsis, downregulation of *ARR7* and *ARR15* causes enlarged meristems (Zhao *et al*., 2010), and overexpression of the Type-C *RR*, *ARR22*, results in dwarf phenotypes with a reduced number of flowers but increased tolerance to cold and drought (Gattolin *et al*., 2006; Kang *et al*., 2013). In rice, *OsRR1* and *OsRR2* positively affect the crown-root formation and negatively affect flowering time (Cho *et al*., 2016; Cho *et al*., 2022; Kitomi *et al*., 2011; Zhao *et al*., 2015; Zhao *et al*., 2009). The double mutant *osrr9 osrr10* exhibits enhanced salt tolerance and slow leaf senescence (Wang *et al*., 2019).

Type-A and -C RR proteins can influence biological processes by interacting with other proteins. Studies have revealed interaction networks between Type-A RRs and other cytokinin signaling components in Arabidopsis and rice. For example, ARR4 interacts with AHP1 in Arabidopsis, and Type-A RRs interact with PHPs and Type-B RRs in rice (Sharan *et al*., 2017; Verma *et al*., 2015). In addition, ARR22, a Type-C RR in Arabidopsis that cannot be induced by cytokinins, interacts with AHP2, AHP3, and AHP5 and is significant in cytokinin signaling (Horak *et al*., 2008). Some Type-A RRs interact with the proteins involved in plant biological rhythms. For example, OsRR1 and OsRR2 interact with Ehd1 to regulate heading date, and ARR4 interacts with phytochrome B to regulate the circadian period (Cho *et al*., 2016; Cho *et al*., 2022; Salome *et al*., 2006; Sweere *et al*., 2001). ARR5 can physically interact with the sucrose nonfermenting1-related kinases SnRK2.2, SnRK2.3, and SnRK2.6, which are significant in the ABA signaling pathway, playing a role in ABA sensitivity and drought tolerance (Huang *et al*., 2018).

Type-A and -C *RRs* significantly affect plant growth by influencing cytokinin signaling; however, studies on *OsRRs* are limited. In this study, we investigated the genetic relationships and expression patterns of Type-A and -C *OsRRs* using RNA-Seq. We evaluated the phenotypes of rice Type-A and -C *OsRR* single- and multiple-mutants produced using CRISPR/Cas9 to determine the functions of different *OsRR*s. Similarly, we analyzed the transcriptomes of young panicles from WT Nipponbare and the *rr2* and *rr4* mutants to identify the genes influenced by RR2 and RR4. Our findings provide new insights into cytokinin signaling and offer valuable references for breeders to discover better germplasm resources.

## Materials and methods

### Plant materials and growth conditions

*Oryza sativa L. ssp. japonica* cultivars Nipponbare (NIP) and Zhonghua 11 (ZH11) were chosen as WT plants for this study. The mutants were *rr1*, *rr2*, *rr3*, *rr4*, *rr5*, *rr7*, *rr9*, *rr10*, *rr9 rr10*, *rr11*, *rr41*, *rr42,* and *rr8 rr12 rr13* in the NIP background and *rr6* in the ZH11 background (Fig. **2**). Double mutant *rr1-2 rr2-2* was a crossing of *rr1-2* and *rr2-2*; *rr4 rr9*, *rr4 rr10,* and *rr4 rr9 rr10* were crossings of *rr4-2* and *rr9 rr10-1*; *rr41-8 rr42-2* was a crossing of *rr41-8* and *rr42-2*.

Rice plants were grown in a field. Field experiments were conducted between 2020 and 2023 in Danyang, Jiangsu Province, China (31.9°N 119.5°E). The sowing dates for the field experiments were May 26, 2020, and May 25 in 2021–2023. Each hill was planted with one seedling with a row spacing of 15 × 30 cm. We used 150 kg/hm^2^ P_2_O_5_ and 240 kg/hm^2^ K_2_O as base fertilizers for the field experiment. And 300 kg ha^−1^ nitrogen (in 2020) or 200 kg ha^−1^ nitrogen (in 2021–2023) with a base fertilizer (to promote seedling growth): tiller fertilizer (to promote tiller growth): panicle fertilizer (to promote panicle development) ratio of 1:1:2. When took photos for WT and mutants, plants were moved from field to pots. For the hydroponic experiment, the nutrient solution comprised 2 mM KNO_3_, 2 mM NH_4_Cl, 0.32 mM KH_2_PO_4_, 0.3 mM MgCl_2_, 0.66 mM CaCl_2_, 0.3 mM Na_2_SiO_3_, 45 μM Fe(II)-EDTA, 11 μM MnCl_2_, 18.5 μM H_3_BO_3_, 0.6 μM Na_2_MoO_4_, 1.4 μM ZnSO_4_ and 1.6 μM CuSO_4_. The nutrient solution was refreshed every four days to maintain a pH of 5.5-6.0.

### Analysis of the panicle structure and seed setting rate

To analyze the panicle structure, we collected the statistics on the number of primary branches, secondary branches, primary spikelets, secondary spikelets, and total spikelets from a minimum of 20 main stem panicles sourced from at least 20 plants. Primary branches refer to those that grow directly on the panicle rachis, whereas secondary branches are those that grow directly on primary branches. Similarly, primary spikelets are directly attached to primary branches, and secondary spikelets grow directly on secondary branches. The lateral organs on the primary branches were considered secondary branches, and the primary spikelets grown on the primary branches were calculated as (secondary branches + primary spikelets)/primary branches. The lateral organs on the secondary branches were spikelets grown on the secondary branches and were calculated as (secondary spikelets)/secondary branches. An illustrative schematic depicting the primary branch, secondary branch, primary spikelet, secondary spikelet, lateral organ on the primary branches, and lateral organ on the secondary branches is shown in Supplemental Figure **S1**.

To assess the seed setting rate, we meticulously documented data from a minimum of 20 main stem panicles across at least 20 plants. The seed setting rate was calculated as the ratio of the number of filled seeds to the total number of spikelets in the entire panicle.

### Plasmids and construction of transgenic plants

To construct the mutants using the CRISPR/Cas9, each *OsRR* was assigned one or two single-guide RNA oligo targets, as previously described (Mao *et al*., 2013). When got the T0 generation of seedling, using PCR for genotyping. For T1 generation, using PCR to screen out the seedlings without T-DNA insertion, and genotyping for further research. The primers used for vector construction and genotyping are listed in Supplementary Table **S1**.

### Sampling, RNA extraction, and gene expression analysis

We sampled young panicles (0–2 mm long, primary and secondary branch differentiation stage) from NIP, *rr2*, and *rr4* plants in the field at the reproductive stage. Each mutant has three biological replicates. The samples were stored at −80 °C until further use. Total RNA was extracted using an E.Z.N.A.^®^ Plant RNA Kit (Omega Biotek Inc.) and subjected to reverse transcription using a PrimeScript RT Reagent Kit (TaKaRa Bio). Quantitative reverse-transcription (qRT)-PCR was performed using a PRISM 7300 Real-Time PCR System (Applied Biosystems) with SYBR^®^ Premix Ex Taq™ (TaKaRa), following the manufacturer’s instructions. *Actin* and *UBQ* were used as internal controls. The primers are listed in Supplementary Table **S7**.

### Sequence alignment and phylogenetic analysis

Amino acid sequence alignment was performed using MUSCLE (Madeira *et al*., 2019), and phylogenetic trees were generated using the neighbor-joining method with 1000 bootstrap iterations in MEGA X (Kumar *et al*., 2018). The protein similarity was calculated using pairwise similarity matrix in TBtools-II (Chen *et al*., 2023).

### RNA-seq and data analysis

To confirm the expression pattern of type-A and type-C RR genes throughout the entire life cycle of rice, we sampled the leaf blade at the vegetative stage (LBV), leaf sheath at the vegetative stage (LSV), **leaf sheath at the reproductive stage (RSV)**, leaf blade at the reproductive stage (LBR), leaf sheath at the reproductive stage (LSR), root at the reproductive stage (RR), stem (S), 0-10mm inflorescence meristem (IM), floret (F), and grains (G). There are two biological replicates for each tissue of leaf blade, leaf sheath, root, stem, floret, and grain. And there are nine biological replicates for the 0-10mm inflorescence meristem (Chang, 2021). To find out differential expressed genes in between NIP and *rr2*, *rr4* mutant, total RNA was extracted from young panicles (0–2 mm long, primary and secondary branch differentiation stage) from NIP, *rr2*, and *rr4* plants using a Trizol reagent kit (Invitrogen, Carlsbad, CA, USA) following the manufacturer’s instructions. Each mutant has three biological replicates. RNA quality was assessed using an Agilent 2100 Bioanalyzer (Agilent Technologies, Palo Alto, CA, USA) and verified through RNase-free agarose gel electrophoresis. After total RNA extraction, the eukaryotic mRNA was enriched with oligo (dT) beads. The enriched mRNA was fragmented into short fragments using fragmentation buffer and reverse-transcribed into cDNA using the NEBNext Ultra RNA Library Prep Kit for Illumina (NEB #7530, New England Biolabs, Ipswich, MA, USA). The purified double-stranded cDNA fragments were end-repaired, tailed with a base, and ligated to Illumina sequencing adapters. The ligation was purified with AMPure XP Beads (1.0X). Subsequently, PCR was performed. The resulting cDNA library was sequenced using an Illumina Novaseq6000 (Gene Denovo Biotechnology Co., Guangzhou, China). The sequencing depth of each sample is 6Gbps.

The obtained reads were processed and analyzed, and genes with False Discovery Rate (FDR) <0.05 and |fold-change| >1.5 were considered to be significantly differentially expressed genes (DEGs). Microsoft Excel was used to create Venn diagrams of DEGs. To gain insight into the changes in panicle phenotype, Gene Ontology (GO) enrichment (http://www.geneontology.org/) and Kyoto Encyclopedia of Genes and Genome (KEGG) pathway (https://www.kegg.jp/) analyses of annotated DEGs were performed using OmicShare tools (http://www.omicshare.com/tools), with the significance level of correction set at *Q*<0.05.

## Results

### Type-A and -C *OsRRs* displayed different expression patterns

The Type-A and -C RR phylogenetic trees comprised five major clades (Fig. **1**), consistent with the findings of a previous study (Tsai *et al*., 2012). *RR3*, *RR5*, *RR6*, *RR7,* and *RR11* were grouped into subfamily A–IV. *RR5* exhibited low expression levels in all tissues surveyed except for the inflorescence meristem. *RR11* shared 82.3% similarity with *RR5* in its amino acid sequence (Supplemental Figure **S2**), showed the highest expression in the inflorescence meristem, and was highly expressed in the leaf sheath and flowers during the reproductive stage. *RR7* was highly expressed in the roots and expressed at lower levels in other organs. *RR3* and *RR6* exhibited higher expression levels than other genes in this subfamily, with *RR3* being highly expressed in the leaf blade during the vegetative stage and in the inflorescence meristem, whereas *RR6* was highly expressed in the roots and flowers.

**Figure 1.**
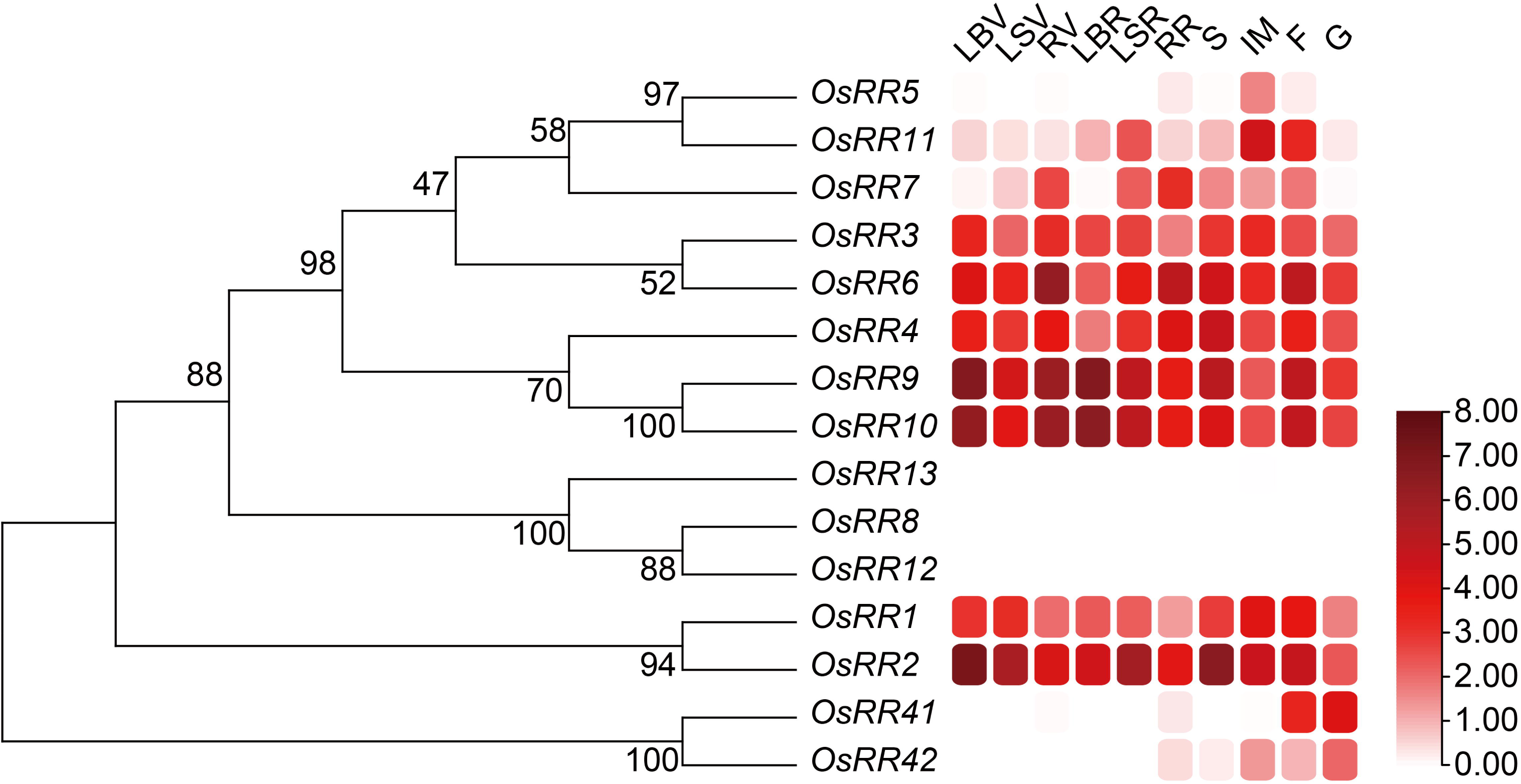
Genetic relationship and expression patterns of Type-A and -C *OsRRs* based on RNA sequencing data. Genetic relationship and expression patterns of Type-A and -C *OsRRs* investigated in the leaf blade at the vegetative stage (LBV), leaf sheath at the vegetative stage (LSV), root at the vegetative stage (RV), leaf blade at the reproductive stage (LBR), leaf sheath at the reproductive stage (LSR), root at the reproductive stage (RR), stem (S), inflorescence meristem (IM), flower (F), and grain (G). The expression patterns were based on log_2_ (Fragments Per Kilobase of exon model per million mapped fragments (FPKM) +1) values.

*RR4*, *RR9*, and *RR10* belonged to the subfamily A-II. They exhibited high expression levels in all plant tissues examined. *RR4* showed the highest expression in the stems. *RR9* and *RR10*, which are homologous genes with 96.1% amino acid sequence similarity, also showed similar expression patterns. Both genes showed the highest expression in the leaf blade during the vegetative and reproductive stages and in the roots during the vegetative stage. The A-V subfamily members *RR8*, *RR12*, and *RR13* shared similar protein sequences (more than 95% compared to each other); however, they exhibited excessively low expression levels in all organs examined.

*RR1* and *RR2* belonged to the A-VI subfamily, and shared the 62.2% similarity. *RR1* exhibited the highest expression in the inflorescence meristem and flowers, whereas *RR2* displayed the highest expression in the leaf blades, leaf sheath, and stem during the vegetative stage.

*RR41* and *RR42* were identified as Type C *RRs*, and both exhibited excessively low expression in vegetative organs. *RR41* showed the highest expression in flowers and grains, while *RR42* had the highest expression in inflorescence meristems and grains.

Each Type-A and -C *RR* was expressed in a distinct pattern. In leaf blades, *RR9* and *RR10* exhibited the highest expression levels at all growth stages, whereas *RR5* and *RR7* showed excessively low expression levels. In the roots, *RR6* showed the highest expression, whereas *RR5* and *RR11* showed markedly low expression levels. In stems, *RR2* was the most highly expressed gene, whereas *RR2* and *RR11* were highly expressed in inflorescence meristems. This suggests that cytokinin signal transduction pathways differ between periods and tissues, enabling plants to finely tune their growth and development. The divergent expression patterns of Type-A and C *RRs* piqued our interest in their functions.

### Most Type-A and C *RR* mutants exhibited reduced plant height and spikelets per panicle

Using CRISPR/Cas9, we constructed single mutants of Type-A and -C *RRs*. Single mutants (except *rr42*) containing at least two mutation sites were successfully generated (Fig. **2a**). Based on their phylogenetic relationships, we also created the *rr9 rr10* double mutant and the *rr8 rr12 rr13* triple mutant (Fig. 2**b**,**c**). All the mutant used in this study showed frameshift mutation (except *RR8* in *rr8 rr12 rr13-2*), the translated proteins of each mutant line were shown in Supplemental Fig. **S3**. To confirm the functions of these genes, we observed their phenotypes throughout their lifecycle.

**Figure 2.**
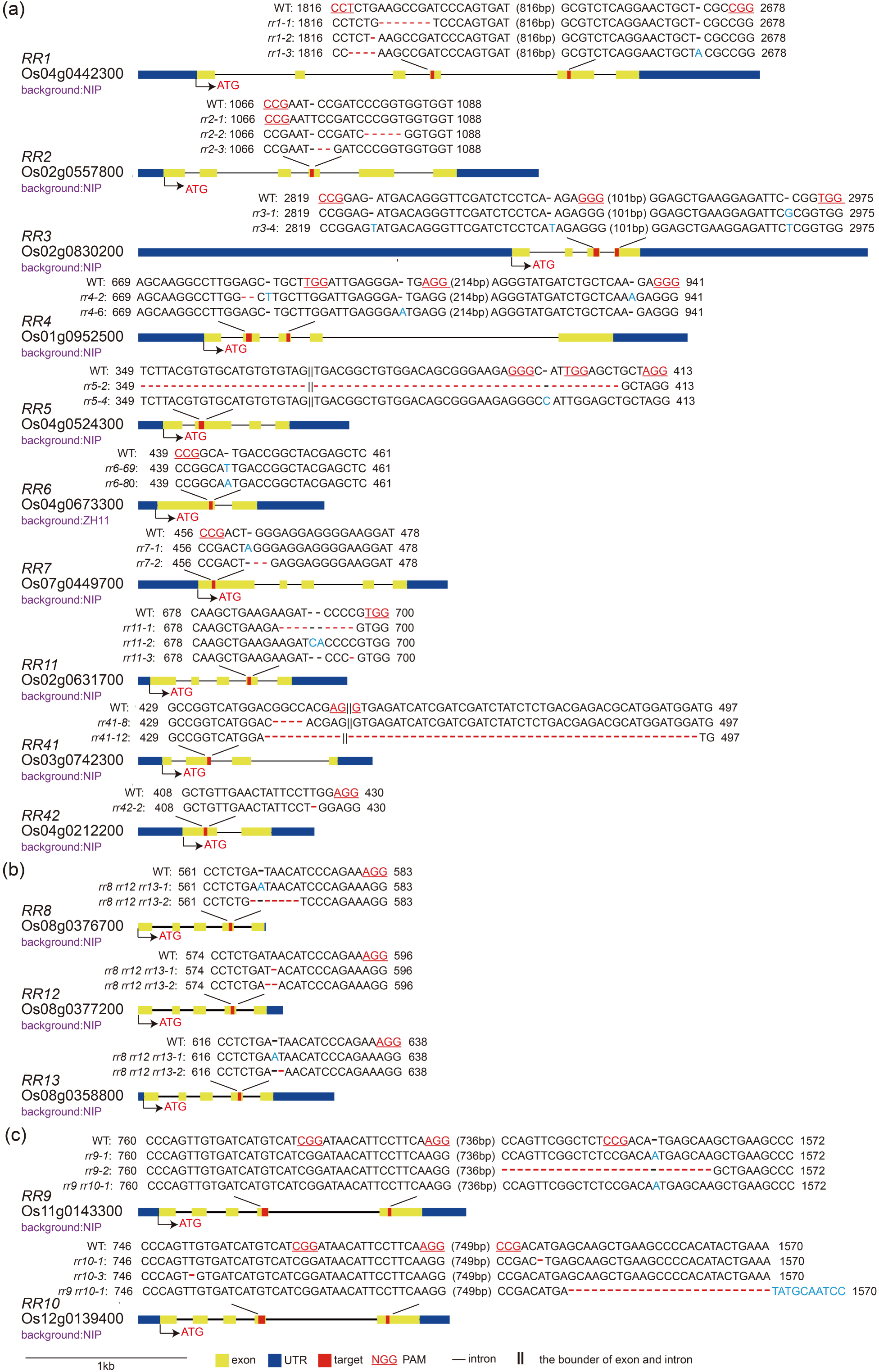
Gene structure and mutation details of single and multiple mutants of Type-A and -C *RRs* in rice. **(a)** Gene structures and mutation details of CRISPR-edited lines targeting *RR1*, *RR2*, *RR3*, *RR4*, *RR5*, *RR6*, *RR7*, *RR11*, *RR41,* and *RR42*. **(b)** Gene structures and mutation details of CRISPR-edited lines in the *RR8, RR12, and RR13* of *rr8 rr12 rr13* mutants. **(C)** Gene structures and mutation details of CRISPR-edited lines in the *RR9 and RR10* of *rr9 rr10* mutants. Solid yellow boxes represent exons, solid lines represent introns, solid blue boxes represent untranslated regions, red boxes represent target sequences, and solid red lines indicate protospacer adjacent motif sequences.

During the vegetative stage, we recorded the number of tillers in all single mutants at the reproductive stage. In 2020, all three *rr1* lines exhibited a significant increase in tiller number, and *rr3-1*, *rr4-6*, and *rr5-4* also exhibited an increased number of tillers (Supplemental Fig. **S4a**). However, the increase in tiller number was slight.

Subsequently, we analyzed the plant heights of the single mutants. The 2-year data showed that most mutants with loss-of-function in Type-A and -C *RRs* exhibited a reduced trend in height, especially *rr7* and *rr10* lines displayed significantly greater reductions across both years (Supplemental Figs. **S4b, S5**). We found that *rr3-1* and *rr4-2* had significantly longer flag leaves (Supplemental Fig. **S4c**). The flag leaves of *rr4-2* and *rr10-3* were wider, whereas *rr7-2* had narrower flag leaves (Supplementary Fig. **S4d**).

Furthermore, we investigated the yield components of the single mutants. As there was no significant change in tiller number initially, the number of panicles in the single mutants also showed no change. Analysis of panicle structure data from the 2 years revealed that *rr5-2*, *rr7-1*, and *rr42-2* exhibited a significant increase in primary spikelets. Most single mutants displayed a decreasing trend in secondary branches and secondary spikelets, particularly several *rr* mutants, such as *rr2*, *rr5*, *rr6*, and *rr10*, where more than one line showed a significant reduction across both years. Owing to the reduced number of secondary spikelets, the total number of spikelets of *rr2*, *rr5*, *rr6*, and *rr10* was also significantly reduced. Moreover, the data from the 2 years indicated that *rr3*, *rr4*, *rr5*, and *rr42* showed reduced seed setting rates compared with that of NIP (Supplemental Tables **S2** and **S3**; Supplemental Fig. **S6**). The loss-of-function of most Type-A and -C *RRs* caused reduced yield factors compared with those of WT, and *rr4-2*, *rr4-6*, *rr5-2*, and *rr42-2* had significantly reduced yields per plant, whereas *rr9 rr10-1* and *rr6-80* showed a significant increase.

### RR1 and RR2 positively regulated plant height and panicle development

Here, we focused on *RR1* and *RR2* functions to elucidate their roles in this subfamily. By hybridizing *rr1-2* and *rr2-2*, we obtained the *rr1-2 rr2-2* (*rr1 rr2*) double mutant to investigate the potential functional overlap or redundancy between the two genes.

Several studies have indicated that RR1 and RR2 regulate root outgrowth (Kitomi *et al*., 2011; Zhao *et al*., 2015; Zhao *et al*., 2009). To confirm their functions during the seedling stage, we grew NIP, three *rr1* lines, two *rr2* lines, and *rr1 rr2* hydroponically for 12 d. At the seedling stage, all single (except *rr2-3*) and double mutants exhibited reduced shoot lengths (Supplemental Fig. **S7a,b**), while *rr1-3*, *rr2-2,* and *rr1 rr2* showed reduced root lengths (Supplemental Fig. **S7c**). However, all mutants showed no change in crown root number (Supplemental Fig. **S7d**).

A previous study reported that higher *RR1* and *RR2* expression negatively affected heading date, as they interacted with Ehd1 (Cho *et al*., 2022). We first examined the heading dates of the NIP, single mutants, and double mutants. We found that the heading date of *rr1-2* was 1.9 d later than that of the NIP, whereas *rr2-2* and *rr1 rr2* had similar heading dates to those of the NIP (Supplemental Fig. **S8a**).

We observed the plant architecture of the NIP, single mutants, and double mutants (Fig. **3a**). In 2022 and 2023, *rr1-2* and *rr2-2* displayed significantly reduced plant height compared with that of NIP, whereas the plant height of the *rr1 rr2* double mutant was shorter than those of the single mutants (Fig. **3b**; Supplemental Table **S4**). Further analysis revealed that the single and double mutants had a shorter top first internode, and the top third and fifth internodes of *rr1 rr2* were reduced (Supplemental Fig. **S8b–d**). However, the internode diameter showed no significant changes in the single or double mutants (Supplemental Fig. **S8e**). Additionally, *rr1 rr2* had shorter and narrower flag leaves, a phenotype that was less pronounced in single mutants (Supplemental Fig. **S8F–H**).

**Figure 3.**
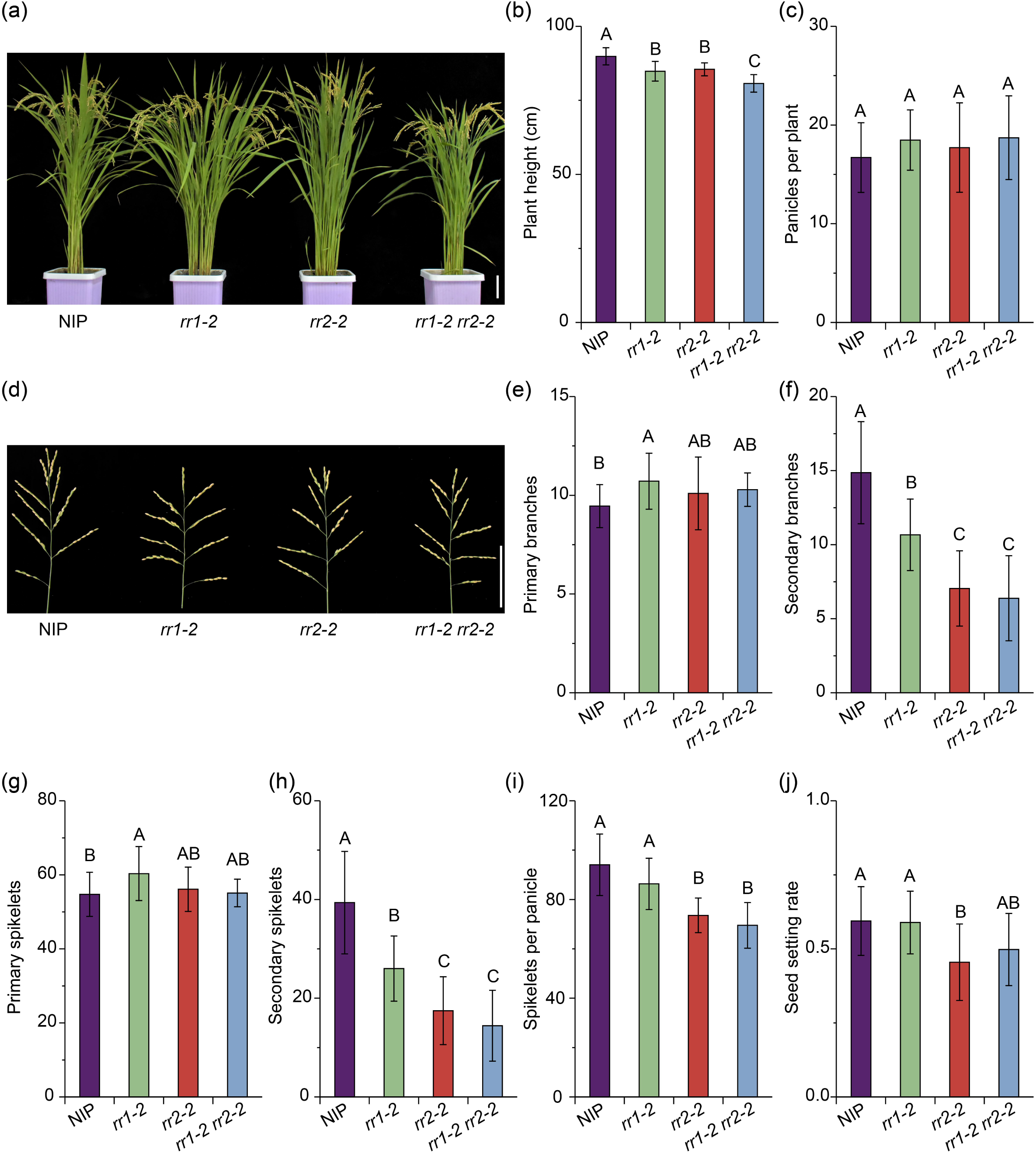
Phenotypic characterization of Nipponbare (NIP) and *rr1*, *rr2*, and *rr1 rr2* mutant plants from the field experiment. **(a)** Whole plant of NIP, *rr1*, *rr2*, and *rr1 rr2*. The image was digitally extracted and scaled for comparison (scale bar = 10 cm). **(b,c)** Measurement of the plant height (**b**) and panicles per plant (**c**) of NIP and *rr1*, *rr2*, and *rr1 rr2* mutants (n>20). Different capital letters indicate the level of statistical significance (*P*<0.01), as determined using Duncan’s multiple range test. **(d)** Panicle phenotype of NIP, *rr1*, *rr2*, and *rr1 rr2*. The image was digitally extracted and scaled for comparison (scale bar = 10 cm). **(e–j)** Measurement of the primary branches per panicle (**e**), secondary branches per panicle (**f**), primary spikelets per panicle (**g**), secondary spikelets per panicle (**h**), spikelets per panicle (**i**), and seed setting rate (**j**) of NIP and *rr1*, *rr2*, and *rr1 rr2* mutants (n>20). Different capital letters indicate the level of statistical significance (*P*<0.01), as determined using Duncan’s multiple range test.

Finally, we focused on the yield structure. The 2-year data showed that the panicle number of *rr1 rr2* was similar to that of the NIP (Fig. **3c**; Supplemental Table **S4**). Regarding the panicle structure, the secondary branches and secondary spikelets of the *rr1 rr2* double mutant were markedly reduced, a phenotype closely resembling that of the *rr2-2* single mutant (Fig. **3d–h**; Supplemental Table **S4**). The total spikelets of *rr1 rr2* were comparable to that of *rr2-2* (Fig. **3i**; Supplemental Table **S4**). To investigate the effects of RR1 and RR2 on panicle development, we calculated the number of lateral organs per primary and secondary branches. The *rr1-2* mutant had significantly fewer lateral organs per primary branch than NIP did, whereas the counts of *rr2-2* and *rr1 rr2* were similar, with an even more pronounced reduction (Supplemental Fig. **S8i**; Supplemental Table **S3**). However, only in 2022, *rr1 rr2* exhibited a significant decrease in lateral organs per secondary branch (Supplemental Fig. **S8j**; Supplemental Table **S4**). Similarly, the 1000-grain weight and seed setting rates of the single and double mutants exhibited varying degrees of reduction compared with those of the NIP, with that of *rr1 rr2* being almost midway between those of *rr1-2* and *rr2-2* (Supplemental Table **S4**).

### In the A-II subfamily, RR4 showed a major effect in regulating panicle development

*RR4*, *RR9*, and *RR10* belonged to the same subfamily. To elucidate the functions of this subfamily, we hybridized *rr4-2* and *rr9 rr10-1*. Subsequently, we obtained the double mutants *rr4 rr9* and *rr4 rr10* and the triple mutant *rr4 rr9 rr10* to analyze their functional overlap or redundancy by observing their phenotypes during the vegetative and reproductive stages in 2022 and 2023 (Supplemental Figure **S9a–c**). In 2022, *rr4-2*, *rr4 rr9*, and *rr4 rr9 rr10* exhibited significantly reduced plant heights and flag-leaf lengths (Supplemental Fig. **S9d,e**). The flag-leaf width of *rr4 rr9 rr10* was also significantly decreased (Supplemental Figure **S9f**). Subsequently, we analyzed their yield components. Notably, in both years, *rr4 rr9*, *rr4 rr10*, and *rr4 rr9 rr10* showed reductions in secondary branches and secondary spikelets, resulting in fewer spikelets per panicle. These phenotypes were similar to those observed in *rr4-2* (Supplemental Table **S5**). The *rr4 rr9*, *rr4 rr10*, and *rr4 rr9 rr10* mutants had lower seed setting rates than the WT did. Furthermore, the 1000-grain weight of *rr4* was lower than that of the WT; however, *rr4 rr9* and *rr4 rr9 rr10* had heavier 1000-grain weights (Supplemental Table **S5**).

### RR8, RR12, and RR13 positively regulated plant height and the development of secondary branches

As summarized above, *RR8*, *RR12*, and *RR13* belonged to the same subfamily and exhibited excessively low expression levels in plant tissues. We also constructed the *rr8 rr12 rr13* triple mutant to determine the functions of this subfamily. For vegetative organs, *rr8 rr12 rr13* displayed reduced plant height, which was particularly noticeable in 2022 (Fig. **4a**; Table **1**). Analysis of the length and diameter of each internode revealed that the length of the top third internode in *the rr8 rr12 rr13* line was significantly reduced, possibly causing the decreased plant height (Supplemental Fig. **S10a,b**). Additionally, in the triple mutants, the diameters of the top first and second internodes were increased (Supplementary Fig. **S10c**). The flag leaf width in *rr8 rr12 rr13* was also decreased (Fig. **4b–d**).

**Figure 4.**
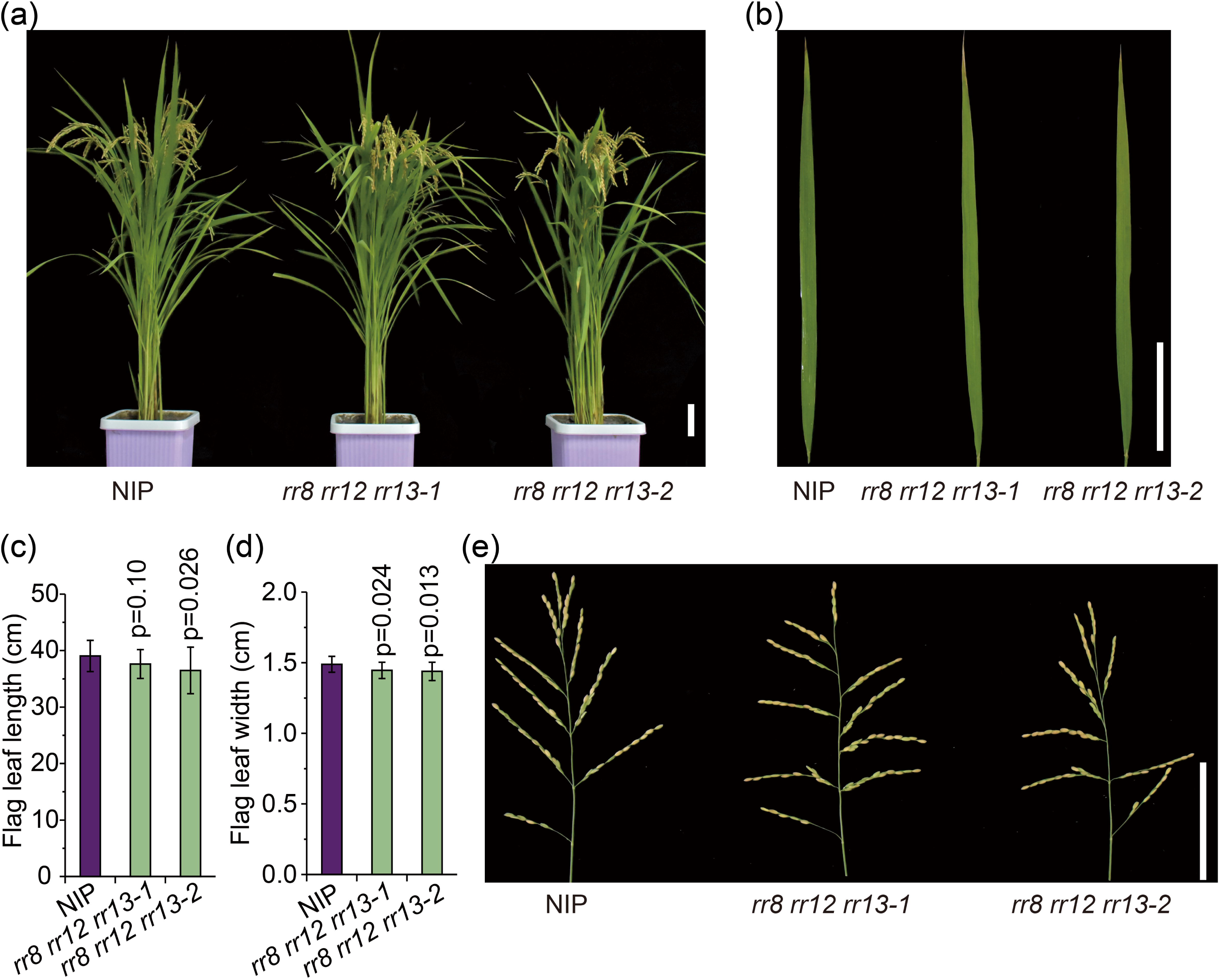
Phenotypic characterization of NIP and *rr8 rr12 rr13* mutant plants in 2022. **(a,b)** Whole plant (**a**) and flag leaf (**b**) phenotypes of NIP and *rr8 rr12 rr13*. The image was digitally extracted and scaled for comparison (scale bars = 10 cm). **(c,d)** measurement of flag leaf length (**c**) and flag leaf width (**d**). Data are shown as means ± SDs (n>10). **(e)** Panicle phenotype of NIP and *rr8 rr12 rr13*. The image was digitally extracted and scaled for comparison (scale bars = 10 cm).

**Table 1.**
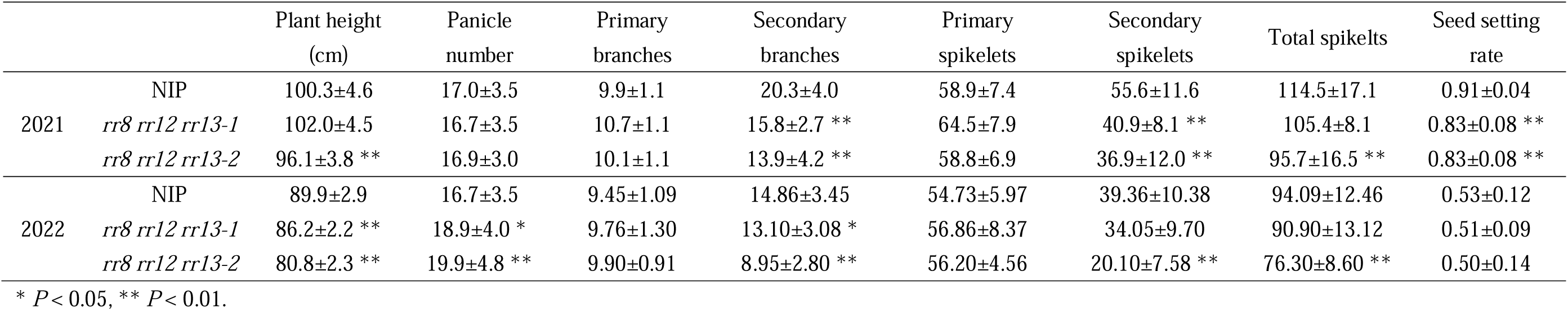
Phenotypes of *rr8 rr12 rr13* mutants.

Subsequently, we examined the yield structure of *rr8 rr12 rr13*. The 2-year data revealed that *rr8 rr12 rr13* exhibited a significant increase in panicle number in 2022 but not in 2021. While both lines of the triple mutant showed no changes in primary branches or primary spikelets, secondary branches were significantly reduced in both years (Fig. **4e**; Table **1**). In addition, the total number of spikelets in the triple mutant showed a decreasing trend. We also observed a decreased seed setting rate in both lines of the *rr8 rr12 rr13* mutant in 2021 (Table **1**), and the *rr8 rr12 rr13-1* line had a higher 1000-grain weight (Supplemental Fig. **S10e**).

### Type-C RR genes function in regulating heading date

Rice contains only two Type-C *RRs*. In a previous study, we investigated the functions of *RR41* and *RR42*. Because they have similar amino acid sequences and expression patterns, whether they exhibit functional redundancy remains unknown. To address this, we crossed *rr41-8* with *rr42-2* to obtain the double mutant *rr41-8 rr42-2*.

First, the loss of function of *RR41* or *RR42* caused earlier heading dates (Supplemental Table **S6**, Supplemental Fig. **S11a,b**). However, *rr41-8 rr42-2* exhibited a heading date similar to that of *rr41-8* and *rr42-2*, indicating that the Type-C *RRs* had no redundancy in regulating the heading date. Moreover, during the vegetative stage, other phenotypes, such as flag leaf length, flag leaf width, plant height, and tiller number, showed no differences in all the single or double mutants compared with those of the WT (Supplemental Table **S6**, Supplemental Fig. **S11c–e**).

Subsequently, we focused on the panicle structure. In 2022, we observed that all single and double mutants had increased numbers of primary branches and primary spikelets compared with those of the WT. Additionally, *rr41-8* exhibited a significant increase in the numbers of secondary branches and secondary spikelets, whereas the numbers were reduced in *rr41-12* and *rr42-2*. Owing to the increased number of primary and secondary spikelets, *rr41-8* displayed a significantly higher spikelet count than the WT did; however, the other mutants showed no significant changes (Supplemental Table **S7**, Supplemental Fig. **S11f**). Furthermore, we observed no significant changes in the number of panicles in any of the mutants. *rr42-2* presented a reduced seed setting rate and 1000-grain weight compared with those of the WT; *rr41-8* and *rr41-12* exhibited an increased 1000-grain weight (Supplemental Table **S7**).

### RNA-seq of gene expression in the young panicle of *rr2* and *rr4*

The data from several years revealed that *RR2* and *RR4* influenced panicle development, particularly by positively regulating secondary branches and secondary spikelets. Therefore, during the reproductive stage, we harvested young panicles from NIP, *rr2*, and *rr4* at the branch differentiation stage and performed RNA-Seq to identify the DEGs (Supplemental Dataset **1**). Compared with the NIP, *rr2* had 637 DEGs (167 upregulated and 470 downregulated) and *rr4* had 594 (130 upregulated and 464 downregulated) (Fig. **5a**). To verify the accuracy and reproducibility of the RNA-seq, we randomly selected eight previously studied genes associated with the phenotypes observed in this study for qRT-PCR. The expression profiles of these genes corroborated the RNA-seq (Supplementary Fig. **S12**).

**Figure 5.**
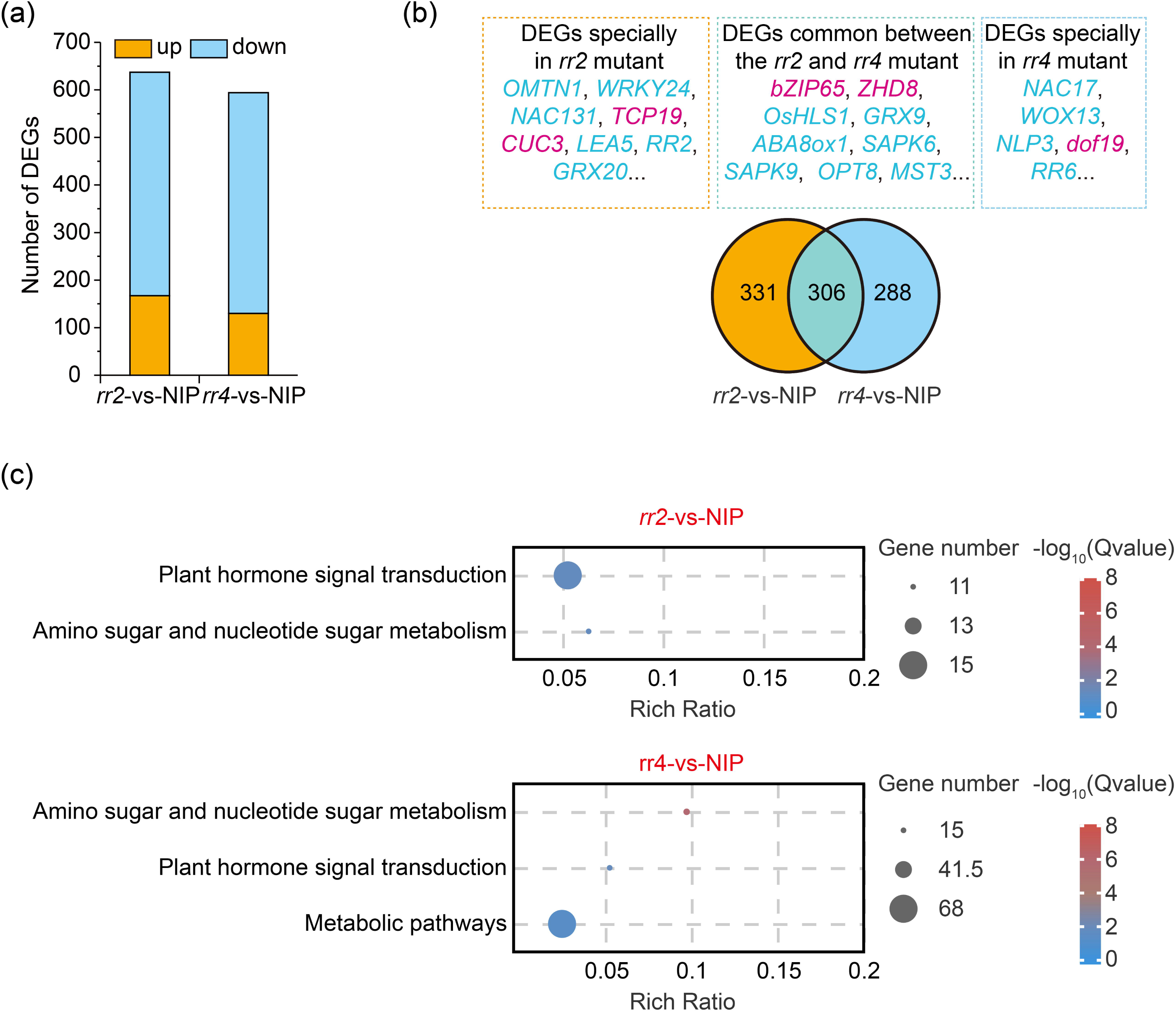
RNA-seq of the young panicles of NIP and *rr2* and *rr4* mutant plants. **(a)** The number of differentially expressed genes (DEGs) identified in the young panicles of the following pairwise comparisons: NIP vs. *rr2* and NIP vs. *rr4*. The changes in gene expression levels were calculated using the log_2_ fold change and *Q* values from three biological replicates. **(b)** Overlapping DEGs in the young panicles of NIP vs. *rr2* and NIP vs. *rr4*. **(c)** Top enriched Kyoto Encyclopedia of Genes and Genome (KEGG) pathways of the DEGs identified in the young panicles of NIP vs. *rr2* and NIP vs. *rr4*. Q values <0.05.

Almost half of the DEGs (306) overlapped between *rr2*-vs-NIP and *rr4*-vs-NIP (Fig. **5b**). The overlapping genes included some transcription factors (such as *bZIP65*, *ZHD8*, and *OsHLS1*), the abscisic acid biodegradation gene *ABA8ox1*, stress-responsive genes (such as *GRX9*, stress-activated protein kinase 6 (*SAPK6)*, and *SAPK9*), and several transporters (such as *OPT8* and *MST3*). In the young panicles of *rr2*, transcription factors, such as *OMTN1*, *WRKY24*, and *NAC131,* were downregulated, whereas *TCP19* and *CUC3* were upregulated. In the young panicle of *rr4*, *NAC17*, *WOX13*, and *NLP3* were downregulated, whereas *dof19* was upregulated (Fig. **5b**). GO and KEGG enrichment analyses were used to extensively examine the DEGs between the mutants and WT. GO analysis revealed that there were 44, 6, and 5 highly enriched GO terms in the biological process, molecular function, and cellular component categories, respectively, in the *rr2* mutant, and 68, 16, and one highly enriched GO terms in the same respective categories in the *rr4* mutant. In *rr2* and *rr4*, the top four enriched GO terms in the biological process category were "Cell wall macromolecule catabolic process," "Response to chemical," "Carbohydrate metabolic process," and "Cell wall macromolecule metabolic process," and the top four enriched terms in the molecular function category were "Chitinase activity," "Hydrolase activity, acting on glycosyl bonds," "Hydrolase activity, hydrolyzing O-glycosyl compounds," and "Chitin binding." In addition, in the cellular component category, both were enriched in the "Extracellular region" (Supplemental Figs. **S13 and S14**). Moreover, in *rr2* and *rr4*, the top enriched KEGG pathways were "Plant hormone signal transduction" and "Amino sugar and nucleotide sugar metabolism" (Fig. **5c**). These suggest that *RR2* and *RR4* are crucial in plant hormone signaling and several other pathways.

### RR2 and RR4 regulated panicle development by influencing several plant hormone pathways

Cytokinins are crucial for panicle development. Type-A *RR*s participate in cytokinin-related pathways and may influence several cytokinin pathways. Regarding cytokinin catabolism, transportation, and signal transduction genes, only *IPT7*, *RR2*, *PUP6,* and *ZOG7* were downregulated in the *rr2* mutant, whereas *RR4*, *RR6,* and *ZOG7* were downregulated in the rr4 mutant. This implies lower cytokinin levels in *rr2* and *rr4* (Fig. **6a**). Heavy metal-associated isoprenylated plant proteins (HIPPs) could inhibit CKX activity by interacting with it, regulating cytokinin homeostasis and signaling (Guo *et al*., 2021). *HIPP56* and *HIPP09* were upregulated in *rr4*, and *HIPP23* and *HIPP39* were downregulated in *rr2* and *rr4* (Fig. 6a), which may influence the cytokinin content.

**Figure 6.**
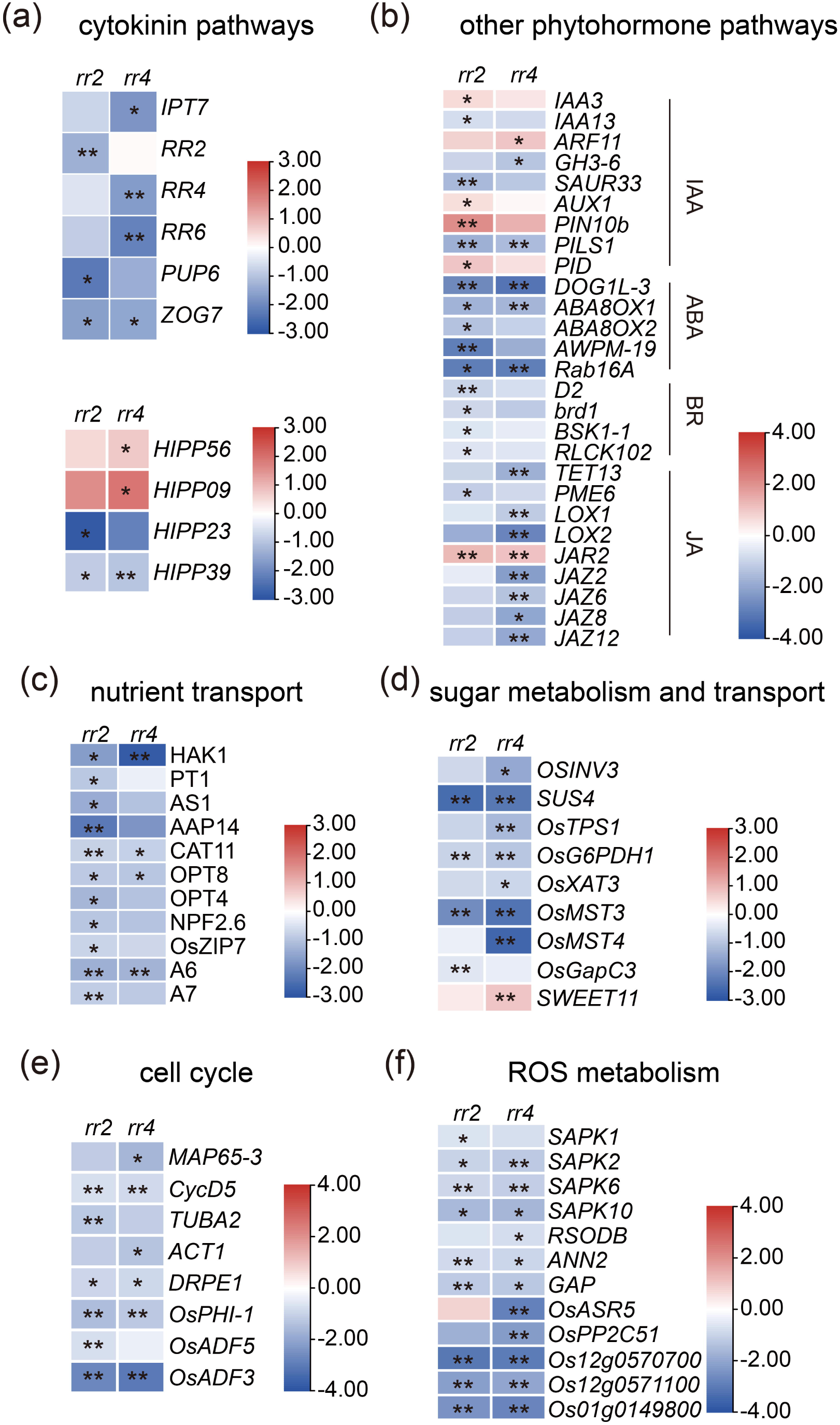
RR2 and RR4 are associated with phytohormone and other pathways. **(a–f)** DEGs related to cytokinin metabolism and signaling transduction (**a**), other phytohormone metabolism and signaling transduction (**b**), nutrition transport (**c**), sugar metabolism and transport (**d**), cell cycle (**e**), and reactive oxygen species (ROS) metabolism (**f**) in the young panicle of different mutants compared with those of NIP. The fold change was based on log_2_ (FPKM_mutant_/FPKM_WT_) values. * *Q*<0.05, ** *Q*<0.01.

Other phytohormones, such as auxin, ABA, and brassinosteroid (BR), are also significant for panicle size. In the auxin pathway, *IAA3*, *IAA13*, *SAUR33*, *AUX1*, *PIN10b*, *PILS1,* and *PID1* were differentially expressed in the young panicles of *rr2*, whereas *ARF11*, *GH3-6,* and *PILS1* were differentially expressed in the young panicles of *rr4* (Fig. **6b**). In the ABA pathway, *DOG1L-3*, which plays a positive role in regulating ABA biosynthesis (Wang *et al*., 2020), was downregulated in *rr2* and *rr4*. Furthermore, the expression of the ABA biodegradation genes *ABA8ox1* and *ABA8ox2*, as well as the ABA-responsive genes *PM-19* and *Rab16A*, displayed a decreasing trend in *rr2* and *rr4*, suggesting that they may have low ABA content in young panicles (Fig. **6b**).

Several BR-related genes, such as *D2*, *brd1*, *BSK1-1*, and *RLCK102*, were significantly downregulated in *rr2*. In *rr4*, jasmonic acid (JA) biosynthesis and signaling genes, including *TET13*, *LOX1*, *LOX2*, *JAZ2*, *JAZ6*, *JAZ8,* and *JAZ12*, were significantly downregulated. These indicate that RR2 and RR4 influence the BR and JA pathways, respectively (Fig. **6b**).

### RR2 and RR4 also influenced other pathways to regulate the panicle structure

To explore additional pathways that RR2 and RR4 may regulate in panicle structure, several essential genes that influence panicle development were examined, such as *SPL7*, *SPL14*, *SPL17*, *LARGE2*, *NLP4*, *APO1*, *APO2*/*RFL*, *LAX1*, *LAX2*, *MOC1*, *MOC3*, *TAW*, *FZP*, *HUB*, *NAM*, *ER1*, *bZIP62*, *DST*, *MADS34*, *MADS5,* and *LP*. However, these genes exhibited no differential expression patterns in the young panicles of *rr2* or *rr4* (Supplemental Fig. **S15**). Several pathways may partly explain the mechanisms by which RR2 and RR4 influence panicle development.

Several genes involved in nutrient assimilation and transport are affected by RR2 and RR4. In *rr2*, the potassium transporter *HAK1*, phosphate transporter *PT1*, nitrogen assimilation gene *AS1*, amino acid/oligopeptide transporter *AAP14*, cation transporter *CAT11*, oligopeptide transporters *OPT4* and *OPT8*, nitrate transporter *NPF2.6*, and iron transporter *ZIP7* were downregulated. This may cause malnutrition in developing panicles, resulting in smaller panicles. Downregulation of *HAK1*, *CAT11*, and *OPT8* was also observed in *rr4*. Plasma membrane H^+^-ATPases participate in nutrient absorption (Loss Sperandio *et al*., 2020; Zhang *et al*., 2021). In this study, two plasma membrane H^+^-ATPases (*A6* and *A7*) showed reduced expression in *rr2*, and *A6* expression was decreased in *rr4* (Fig. **6c**). Sugar-related genes were also influenced by RR2 and RR4. Several sugar-related genes, including the invertase gene *INV3*, sucrose synthase gene *SUS4*, sugar transporters such as *MST3* and *MST4*, and metabolism genes such as *G6PDH1* and *GAPC3*, were downregulated in *rr2* and *rr4*, which could impact panicle development (Fig. **6d**).

Second, in *rr2* and *rr4*, several microtubule, tubulin, and cell phase genes were downregulated, which could cause the formation of smaller panicles owing to decreased cell numbers in the young panicle (Fig. **6e**).

Finally, in *rr2* and *rr4*, several SAPK genes that could help clear reactive oxygen species (ROS) were downregulated in young panicles. Other genes involved in ROS metabolism were also influenced by RR2 and RR4. In the young panicles of *rr2* and *rr4*, *GAP* and *ANN1* were downregulated, while *ASR5* was downregulated in *rr4*. A copper/zinc superoxide dismutase, *RSODB*, was also downregulated in *rr4*. Three metallothioneins (Os12g0570700, Os12g0571100, and Os01g0149800) were also downregulated in the *rr2* and *rr4* mutants (Fig. **6f**). These suggest that *rr2* and *rr4* exhibit weaker ROS metabolic activity and that high ROS accumulation may also inhibit the differentiation of branches and spikelets.

## Discussion

### Type-A and -C *RRs* showed little functional differences

Cytokinins play a positive role in promoting tiller bud outgrowth and panicle development, contributing to increased yield. Researchers have suggested that Type A and C RR proteins with similar structures can inhibit cytokinin signal transduction (Horak *et al*., 2008; Tsai *et al*., 2012). However, most studies use the expression level of type-A and type-C *RR* genes to explain cytokinin content in plant tissues and phenotypes. Yet, there is a lack of studies explaining the function of type-A and type-C *RR* genes. In this study, we systematically identify the expression pattern and function of these genes.

The transcriptome data showed that several Type-A *RRs* (except *RR5*, *RR8*, *RR12*, and *RR13*) were highly expressed in various plant tissues, particularly in young panicles. Type-C *RRs* were specifically expressed during the reproductive stage. Moreover, Type A RR proteins were primarily localized in the cytosol and nucleus, and interactions between Type A and B RR proteins occurred in the nucleus (Sharan *et al*., 2017; Tsai *et al*., 2012). Based on these findings, we hypothesized that Type-A and -C *RRs* show specific expression patterns that could help regulate cytokinin signaling in various plant tissues and cell organs. However, owing to past limitations in genome editing technology, studies on the function of Type A and C *RRs* have some shortcomings. Several studies have indicated that *RR1* and *RR2* influence crown root outgrowth in rice (Kitomi *et al*., 2011; Zhao *et al*., 2015; Zhao *et al*., 2009). However, our study showed that the *rr1* and *rr2* single mutants and the *rr1 rr2* double mutant showed a similar number of crown roots to that of the WT. As several type-A *RR* genes (such as *RR1*, *RR2*, *RR3*, *RR4*, *RR6*, *RR7*, *RR9* and *RR10*) showed highly expression in root, suggesting that more Type-A *RRs* may commonly regulate crown root outgrowth. Moreover, adding to previous research, we focused on shoot development in *rr* mutants. After summarizing the phenotypes of some Type-A and -C *rr* mutants over several years, we discovered some findings. For vegetative organs, we observed no consistently different flag-leaf lengths and widths among the single mutants. In contrast, we found that *RR1* and *RR2* exhibited functional redundancy in flag-leaf development; *rr1* and *rr2* exhibited no noticeable differences in flag-leaf lengths and widths; however, the *rr1 rr2* double mutant showed significantly shorter and narrower flag leaves than the WT and single mutants did. Most Type-A and -C *RRs* play subtle positive roles in plant height. They may have a functional overlap, as the *rr1 rr2* double mutant was shorter than the *rr1* or *rr2* single mutant.

Ehd1 promotes the transition from the vegetative to reproductive stage in rice by inducing the expression of *FT-like* genes, regulating the heading date (Takahashi *et al*., 2009). RR1 and RR2 interact with Ehd1 and disrupt its function by inhibiting homodimerization (Cho *et al*., 2016; Cho *et al*., 2022). *RR1* or *RR2* overexpression in rice resulted in delayed heading dates, suggesting that *RR1* and *RR2* play negative roles in flowering time. Hence, we focused on the heading dates of *rr1*, *rr2,* and *rr1 rr2*. The *rr1 rr2* double mutant exhibited a heading date similar to that of the WT. This suggests that other Type-A *RRs* may also have functional redundancy in regulating heading date. Additionally, we found that Type-C *RRs* had negative effects on the heading date. However, the mechanisms by which *RR41* and *RR42* modulate the floral transition remain unknown.

Regarding yield factors, almost no Type A or -C *RRs* influenced the number of panicles. However, several Type-A *RRs*, including *RR1*, *RR2*, *RR4*, *RR5*, *RR6*, *RR7*, *RR8*, *RR10*, *RR12*, and *RR13*, played positive roles in panicle development, particularly in increasing the number of secondary branches and spikelets, leading to an increase in total spikelets. This is highly related to the higher expression of type-A *RR* genes in young panicles. Contrary to our expectation that each branch of the *RR* gene subfamily in the evolutionary tree would have functional redundancy, each branch contained a major gene regulating the panicle structure. For example, the secondary branches and spikelets of *rr1 rr2* were similar to those of *rr2*, whereas those of *rr4 rr9*, *rr4 rr10*, and *rr4 rr9 rr10* were similar to those of *rr4*. Furthermore, *RR3*, *RR5*, and *RR42* positively regulated the seed setting rate, and some Type-A and -C *RRs* affected the 1000-grain weight.

In contrast to the functional divergence observed in *CKXs*, loss-of-function mutations in Type-A and -C *RRs* caused fewer phenotypic effects. Based on the analysis of mutant phenotypes in this study, we suggest that Type-A and -C *RRs* have similar roles in panicle development and plant height (Fig. **7a**); however, they exhibit slight differences in regulating grain weight. However, their phenotypes were not evident, suggesting that this group of genes exhibits functional overlap or redundancy.

**Figure 7.**
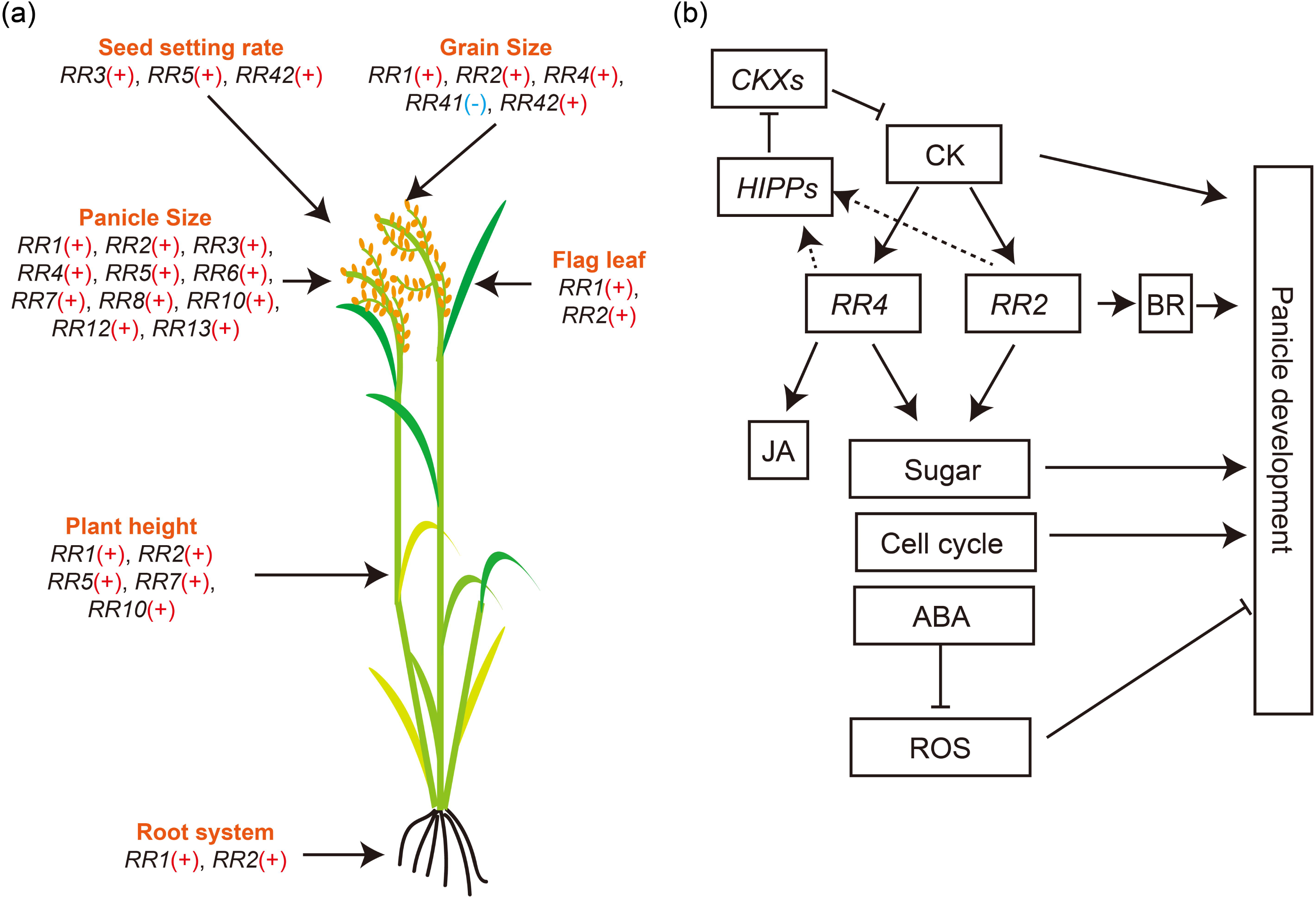
Functions of Type-A and -C *RRs* in rice. (**a**) The function of each Type-A and -C *RRs* in regulating the growth and development of each organ in rice summarized from the phenotypes of all mutants. “+” means positive regulation, while “-” means negative regulation. (**b**) The regulation network of RR2 and RR4 during rice panicle development.

### RR2 and RR4 positively regulated panicle development through several pathways

Our phenotypic studies revealed that almost all Type-A *RRs* participated in panicle development. As *rr2* and *rr4* exerted major effects within their respective subfamilies, we performed RNA-seq on young panicles to elucidate the pathways through which RR2 and RR4 regulate panicle development. After analyzing the data, we constructed a control network for RR2 and RR4 functions in young panicles (Fig. **7b**).

First, a feedback loop may exist between cytokinin signaling and accumulation. The expression levels of Type-A *RRs* were downregulated in the young panicles of *rr2* and *rr4*, suggesting that Type-A *RRs* may positively regulate cytokinin accumulation. A previous study reported that *OsRR6*-overexpression lines exhibited lower *CKX* expression and increased cytokinin content (Hirose *et al*., 2007). These imply that RR2 and RR4 may upregulate cytokinin content in young panicles, promoting panicle differentiation. RR2 and RR4 may influence the cytokinin content via the HIPP-CKX pathway.

RR2 and RR4 also influence other plant hormone-related pathways. ABA biosynthesis is commonly induced by RR2 and RR4. RR2 and RR4 induced BR and JA biosyntheses, respectively. Previous studies have shown that some Type-A *RRs*, such as *RR7*, are upregulated by auxins (Tsai *et al*., 2012). This suggests that Type-A *RR* genes may have functional differences and serve as bridges for plant hormone interactions.

In addition, we found that RR2 and RR4 promoted sugar metabolism, transport, and cell division, which may promote panicle development. ABA reduced the ROS content, benefiting panicle development. Recent studies have shown that high ROS levels can cause spikelet degeneration (Zhang *et al*., 2024). A moderate amount of ABA could support ROS metabolism and help spikelets reduce oxidative damage, whereas low ABA content (as in *nced5* mutants) could be detrimental to spikelet development owing to increased ROS levels in young panicles. High BR levels can enhance ROS metabolism (Zhang *et al*., 2024). In *rr2* and *rr4*, low ABA levels caused ROS accumulation, facilitated by a reduction in ROS-scavenging genes. Additionally, SAPK6, which acts downstream of ABA and is downregulated in the young panicles of *rr2* and *rr4*, phosphorylates SPL14 and increases its stability at the seedling stage (Jia *et al*., 2022). As SPL14 can enlarge the panicle size, a similar process regulated by RR2 and RR4 may also occur in young panicles, increasing the panicle size.

Our systematic analysis of 13 Type-A *RRs* and two Type-C *RR* genes in rice revealed their complex expression patterns. We generated 34 *RR* mutant lines using CRISPR/Cas9 or hybridization and observed the phenotypes of the mutants throughout the growth period. This enabled the determination of the functions of specific genes involved in plant development. We found that RR2 and RR4 had the strongest function in promoting panicle development. Using RNA-seq, we identified several pathways influenced by RR2 and RR4 and found that the regulatory pathways shared functional divergence and overlap. Our findings provide a significant resource for systemically elucidating the function of Type-A and -C RR genes and provide new insights for defining cytokinin signal transduction in future studies.

## Acknowledgments

We thank Jiankang Zhu and Caixia Gao for providing the vectors of the CRISPR-Cas9 system. We also thank Biogle and Biorun genome editing center for producing transgenic rice. This work was supported by the Natural Science Foundation of Jiangsu Province (Grants No BK20231470).

## Author contributions

C.D. and Y.D. conceived the original screening and research plans; C.R., R.Z., J.X., T.Y., Z.L., Y.L., R.X., X.S., J.L., X.Z., J.S., Y.M., Z.C., and C.D. performed the experiments and analyzed the data; C.R. and C.D. wrote the article with contributions of all the authors; C.D. agrees to serve as the author responsible for contact and ensures communication.

## Data availability

All data supporting the findings of this study are available within the paper and within its supplementary data published online.

## Supplemental data

The following supplemental materials are available.

**Supplemental Figure S1.** Schematic representation of rice inflorescences

A primary branch growing on the rachis is indicated by a lavender circle, and a secondary branch growing on the primary branch is indicated by a light green circle. Spikelets growing directly on primary branches are referred to as primary spikelets (magenta), whereas those growing on secondary branches are referred to as secondary spikelets (azure). The lateral organs on the primary branches represent the secondary branches, and the primary spikelets grow on the primary branches, indicated by orange arrows. The lateral organs on the secondary branches represent secondary spikelets grown on the secondary branches, as indicated by the orange arrows.

**Supplemental Figure S2.** Similarity Matrix of type-A and -C Proteins in rice

**Supplemental Figure S3.** Protein mutation details of single and multiple mutants of Type-A and -C RRs in rice

Gene structures and mutation details of CRISPR-edited lines targeting *RR1*, *RR2*, *RR3*, *RR4*, *RR5*, *RR6*, *RR7*, *RR11*, *RR41*, and *RR42* in single mutants, *RR8*, *RR12*, and *RR13* of *rr8 rr12 rr13* mutants, *RR9* and *RR10* of *rr9 rr10* mutants. Solid yellow boxes represent amino acid sequence, red boxes represent mutation sites, and amino acid in black represents amino acid without change compared to WT, in red represents amino acid changed compared to WT.

**Supplemental Figure S2.** Phenotypic characterization of vegetative organs in the *rr* mutants from the 2020 field experiment

**a–d**: Measurement of tiller number at the vegetative stage 70 days after sowing (**a**), plant height (**b**), flag leaf length (**c**), and flag leaf width (**d**) between single mutants and their wild-type background (n>20 for **a–d**). *P*-values indicate the level of statistical significance between wild-type and single mutants, as determined using Student’s *t*-test.

**Supplemental Figure S3.** Phenotypic characterization of plant height in the *rr* mutants from the 2021 field experiment

Comparison of plant height between single mutants and their wild-type background (n>20). *P*-values indicate the level of statistical significance between wild-type and single mutants, as determined using Student’s *t*-test.

**Supplemental Figure S4.** Images of the *rr* mutant panicle structure of NIP from the 2021 field experiment

Scale bars = 10 cm.

**Supplemental Figure S5.** Phenotypic characterization of Nipponbare wild type, *rr1*, *rr2*, and *rr1 rr2* seedlings

**a:** Phenotypic features of whole plants at the seedling stage of Nipponbare (NIP), *rr1*, *rr2,* and *rr1 rr2* plants. The images were digitally extracted and scaled for comparison. Scale bar = 10 cm. **b–d**: Measurement of shoot length (**b**), root length (**c**), and crown root number (**d**); data are shown as means ± SDs (n>20). Different capital letters indicate the level of statistical significance (*P*<0.01), as determined using Duncan’s multiple range test.

**Supplemental Figure S6.** Phenotypic characterization of Nipponbare wild-type, *rr1*, *rr2,* and *rr1 rr2* **a**: Measurement of the heading date of the NIP, *rr1*, *rr2,* and *rr1 rr2*. Data are shown as means ± SDs (n>40). *P*-values indicate the level of statistical significance between wild-type and single mutants, determined using Student’s *t*-test. **b,c:** Main stems (**b**), and internode and panicle length phenotypes (**c**) of NIP, *rr1*, *rr2,* and *rr1 rr2*. The images were digitally extracted and scaled for comparison. Scale bar = 10 cm. **d:** Measurement of the lengths of each internode and panicle in NIP, *rr1*, *rr2,* and *rr1 rr2* (n>20). Different capital letters indicate the level of statistical significance (*P*<0.01), as determined using Duncan’s multiple range test. **e**: Measurement of the diameter of each internode in NIP, *rr1*, *rr2,* and *rr1 rr2* (n>20). Statistical significance is indicated by \**P*<0.05 and \*\**P* < 0.01. F, Flag leaf phenotypes of NIP, *rr1*, *rr2,* and *rr1 rr2* mutants. The images were digitally extracted and scaled for comparison. Scale bar = 10 cm. **g–j**: Measurement of flag leaf length (**g**), flag leaf width (**h**), lateral organs on primary branches (**i**), and lateral organs on secondary branches (**j**). Data are shown as means ± SDs (n>20). Different capital letters indicate the level of statistical significance (*P*<0.01), as determined using Duncan’s multiple range test.

**Supplemental Figure S7.** Phenotypic characterization of Nipponbare wild type, *rr4-2*, *rr9 rr10-1*, *rr4 rr9*, *rr4 rr10,* and *rr4 rr9 rr10*

**a–c**: Phenotypic features of whole plants at the mature stage (**a**), panicle phenotype (**b**), and flag leaf phenotype (**c**) of Nipponbare (NIP), *rr4-2*, *rr9 rr10-1,* and *rr4 rr9 rr10* plants. The images were digitally extracted and scaled for comparison. Scale bar = 10 cm. **d–h**: Measurement of plant height (**d**), flag leaf length (**e**), and flag leaf width (**f**); data are shown as means ± SDs (n>10). Different capital letters indicate the level of statistical significance (*P*<0.01) as determined using Duncan’s multiple range test.

**Supplemental Figure S8.** Phenotypic characterization of Nipponbare wild-type and *rr8 rr12 rr13* mutant

**a,b**: Main stem (**a**) and internode and panicle length (**b**) phenotypes of NIP and *rr8 rr12 rr13* mutant. The images were digitally extracted and scaled for comparison. Scale bars = 10 cm. **c,d**: Measurement of the length (**c**) and diameter (**d**) of each internode and panicle in the NIP and *rr8 rr12 rr13* mutant (n>20). Statistical significance is indicated by \**P*<0.05 and \*\**P*<0.01. **e**: Measurement of the 1000-grain weight. Data are shown as means ± SDs (n>10). *P*-values indicate the level of statistical significance between wild-type and single mutants determined using Student’s *t*-test.

**Supplemental Figure S9.** Phenotypic characterization of Nipponbare wild-type and *rr41*, *rr42,* and *rr41 rr42* mutants

**a,b**: Phenotypic features of whole plants at the heading date of NIP, *rr41-8* and *rr41-12* (**a**); NIP, *rr41-8*, *rr42-2*, and *rr41 rr42* (**b**). **c,d**: Phenotypic features of whole plants at the mature stage of NIP, *rr41-8,* and *rr41-12* (**c**); NIP, *rr41-8*, *rr42-2*, and *rr41 rr42* (**d**). **e,f**: Phenotypic features of flag leaves (**e**) and panicle structure (**f**) of NIP, *rr41-8*, *rr41-12*, *rr42-2,* and *rr41 rr42*. The images were digitally extracted and scaled for comparison. Scale bar = 10 cm.

**Supplemental Figure S10.** FPKMs and relative expression levels of *OMTN1*, *PIN10b*, *TCP19*, *AS1*, *PUP6*, *ZOG7*, *RR2,* and *RR4* in the young panicles of Nipponbare (NIP), *rr2*, and *rr4*. Quantitative expression changes in eight differentially expressed genes using RNA-seq and qRT-PCR. The results are presented relative to those of the NIP. The FPKM value and qRT-PCR data represent means ± SD (n = 3). Total RNA was isolated from NIP or mutant young panicles. β*-Actin* and *Ubq* were used as reference genes.

**Supplemental Figure S11.** Significantly enriched GO terms in the three categories (biological processes, molecular functions, and cellular components) between NIP and *rr2*

**Supplemental Figure S12.** Significantly enriched GO terms in the three categories (biological processes, molecular functions, and cellular components) between NIP and *rr4*

**Supplemental Figure S13.** Differentially expressed genes related to panicle development in the young panicles of Nipponbare wild type and *rr2* and *rr4* mutants Differentially expressed genes related to panicle development in the young panicles of different mutants compared with that of Nipponbare (NIP). The fold-change was based on log_2_ (FPKM_mutant_/FPKM_WT_) values. * *Q*<0.05, ** *Q*<0.01.

**Supplemental Table S1.** Information on the primers used in the study

**Supplemental Table S2.** Yield structure of *rr* mutants in 2020

**Supplemental Table S3.** Yield structure of *rr* mutants in 2021

**Supplemental Table S4** Phenotypes of NIP, *rr1*, *rr2* and *rr1 rr2* in 2022

**Supplemental Table S5.** Yield phenotypes of NIP, *rr4-2*, *rr9 rr10-1*, *rr4 rr9*, *rr4 rr10* and *rr4 rr9 rr10*

**Supplemental Table S6.** Phenotypes of Type-C *RR* mutant*s* in the vegetative stage in 2022

**Supplemental Table S7.** Yield structure of Type-C *RR* mutants in 2022

**Supplemental Dataset 1.** RNA sequencing data

